# Glutathione-S-transferase from the arsenic hyperaccumulator fern *Pteris vittata* can confer increased arsenate resistance in *Escherichia coli*

**DOI:** 10.1101/379511

**Authors:** Aftab A. Khan, Danielle R. Ellis, Xinyuan Huang, Gareth J. Norton, Andrew A. Meharg, David E. Salt, Laszlo N. Csonka

## Abstract

Although arsenic is generally a toxic compound, there are a number of ferns in the genus *Pteris* that can tolerate large concentrations of this metalloid. In order to probe the mechanisms of arsenic hyperaccumulation, we expressed a *Pteris vittata* cDNA library in an *Escherichia coli ΔarsC* (arsenate reductase) mutant. We obtained three independent clones that conferred increased arsenate resistance on this host. DNA sequence analysis indicated that these clones specify proteins that have a high sequence similarity to the phi class of glutathione-S-transferases (GSTs) of higher plants. Detoxification of arsenate by the *P. vittata* GSTs in *E. coli* was abrogated by a *gshA* mutation, which blocks the synthesis of glutathione, and by a *gor* mutation, which inactivates glutathione reductase. Direct measurements of the speciation of arsenic in culture media of the *E. coli* strains expressing the *P. vittata* GSTs indicated that these proteins facilitate the reduction of arsenate. Our observations suggest that the detoxification of arsenate by the *P. vittata* GSTs involves reduction of As(V) to As(III) by glutathione or a related sulfhydro compound.

**Funding:** The authors acknowledge support from the Indiana 21st Century Research and technology Fund (912010479) to DES and LNC, from the U.S. Department of Energy (grant no. DE-FG02-03ER63622) to DES, and from BBSRC-DFID (grant no. BBF0041841GJN) to AAM. The funders had no role in study design, data collection and analysis, decision to publish, or preparation of the manuscript. There are no financial, personal, or professional interests that could be construed to have influenced the paper.

## Introduction

Several members of the fern genus *Pteris* can accumulate high levels of arsenic [1,2]. The first arsenic hyperaccumulator discovered was *Pteris vittata* (Chinese brake fern), which can contain arsenic at levels in excess of 20 mg/g in its fronds [1,3]. Such high level accumulation, and hence, tolerance of arsenic is remarkable in view of the toxicity of this element [4]. The exceptionally high arsenic tolerance of the hyperaccumulating ferns not only makes them good candidates for use in biological remediation of arsenic-contaminated environments, but it also makes them interesting systems to unravel the basic mechanisms of resistance to this widespread environmental pollutant.

Paradoxically, detoxification of arsenate involves reduction to arsenite [5,6], which has potentially higher human and ecological toxicity than arsenate [7,8]. Arsenite produced from arsenate is known to be removed from the cytoplasm by various efflux mechanisms. In *Escherichia coli,* it is pumped out by an ATP-dependent export system, and in yeast it is eliminated by a combination of efflux into the medium by a plasma membrane-bound carrier and transfer into the vacuole as a glutathione adduct [5,6]. In *Arabidopsis thaliana,* an arsenic sensitive non-hyperaccumulating plant, arsenite is transferred to the vacuole, as a phytochelatin adduct via two paralogous ABCC-type transporters *(AtABCC1* and *AtABCC2)* [9]. Three different families of arsenate reductases have been identified, represented by the ArsC protein of *E. coli* [10], the ArsC protein of *Staphylococcus aureus* [11], and the Acr2 protein of yeast [12]. The *E. coli* and yeast arsenate reductases use reduced glutathione (GSH) as the direct electron donor for the reduction of arsenate, whereas the *S. aureus* enzyme employs thioredoxin. Ellis et al. [13] cloned the *P. vittata PvACR2* arsenate reductase gene from a cDNA library of arsenate-treated gametophytes. This enzyme shows 25% amino acid sequence identity / 47% similarity to the yeast ACR2 protein and likewise uses GSH as substrate. Like the yeast enzyme, the *P. vittata* ACR2 lacks phosphatase activity [13]. Arsenate reductase has been shown to be induced 7-fold by arsenate in the roots of *P. vittata* [14]. The reduction product of arsenate is accumulated as free arsenite in the fronds of this fern probably in the vacuoles [3,13,15]. Arsenate reductases similar to the *P. vittata* ACR2 protein have been cloned from rice [16] and the arsenate-tolerant grass *Holcus lanatus* [4], and orthologous sequences have been identified in the genomes of other plants and fungi that are not arsenic hyperaccumulators [16], though all these enzymes also have phosphatase activity.

Phytochelatins are cysteine-rich peptides derived from GSH that can bind to arsenite and several reactive heavy metal cations [17]. *A. thaliana* mutants unable to synthesize phytochelatins show enhanced sensitivity to arsenic [18], and overproduction of these compounds has been shown to increase arsenate resistance [4,19,20]. Phytochelatins have been detected in *P. vittata* and in the related species *Pteris cretica,* but unlike in non-hyperaccumulator plants, only a small portion of arsenic is complexed with phytochelatins in these hyperaccumulators. Phytochelatin production in these arsenic hyperaccumulators is limited [17,21], suggesting that arsenic-phytochelatin complexes do not form a major storage form for arsenic, although they still may play an important role in hyperaccumulation, as indicated by the occurrence of highly localized thiolate-coordination of arsenic in close proximity to the vein and mid-veins in the fronds of the *P. vittata* sporophyte [17,21].

Two other proteins that could play a role in arsenic tolerance or hyerpaccumulation in *P. vittata* have been identified by selection for clones from cDNA libraries that increased the arsenate tolerance of an arsenate reductase mutant *(ΔarsC)* of *E. coli.* One gene obtained by this approach was found to specify a triosephosphate isomerase [22]. In addition to its expected enzymatic activity, it was suggested that this enzyme promotes the reduction of arsenate by an unknown mechanism. The second enzyme that was identified in this manner was a glutaredoxin [23], which was able to confer increased resistance to both arsenate and arsenite by a mechanism that needs to be determined.

In order to identify other proteins that could contribute to the arsenic resistance of *P. vittata*, we carried out a similar procedure to isolate clones from a *P. vittata* cDNA library that elevated the arsenate resistance of a *ΔarsC* mutant *E. coli.* Nucleotide sequence analysis revealed that the cDNA clones in these derivatives encode proteins that have high sequence similarity to glutathione-S-transferases (GSTs).

## Materials and Methods

### Bacterial Strains and Growth Conditions

The *E. coli* K12 strain W3110 (IN[rrnD-rrnE]) and its arsenate reductase mutant derivative WC3110 *(ΔarsC IN[rrnD-rrnE])* [12] were obtained from B. Rosen (Wayne State University), the γ-glutamyl-cysteine synthetase mutant JTG10 *(gshA20::Tn10kan thr-1 araC14 leuB6[Am] Δ(gpt-proA)62 lacY1 tsx-33 glnV44galK2 hisG4 rfbC1 mgl-5 kdgK51 xylA5 mtl-1 argE3 thi-1 λ rpsL31)* [24] was obtained from the *E. coli* Genetic Stock Center, and the glutathione reductase mutant BW31342 *(Δgor::cat)* and the arsenate reductase mutant BW39212 *(ΔarsC::kan)* were obtained from B. Wanner (Purdue University). The arsenate reductase-glutathione synthetase double mutant strain KC1680 was constructed by P1 transduction of the gshA20::Tn10kan from strain JTG10 into WC3110. The arsenate reductase-glutathione reductase double mutant strain KC1852 was constructed by transduction of the *ΔarsC::kan* and *Δgor::cat* insertions elements BW31342 and BW39212 into WC3110. Strains KC1858 *(Δgor::cat ΔarsC::kan IN[rrnD-rrnE]* / pTriplEx2), KC1860 *(ΔarsC::kan Δgor::cat* IN[rrnD-rrnE] / *pPvGST35),* and KC1862 *(ΔarsC::kan Δgor::cat* IN[rrnD-rrnE] / *pPvGST41)* were made by electroporating the indicated plasmids into KC1852.

Bacterial cultures were grown with aeration at 37^°^ in the rich medium LB [25] or K medium, which has a phosphate concentration of 1 mM [26]. When used, ampicillin (Ap), chloramphenciol (Cm), and kanamycin sulfate (Km) were at 100, 25, and 75 mg l^-1^, respectively. Solid media contained 20 g l^-1^ Difco agar. Na_2_AsO_4_·7H_2_O and NaAsO_2_ were purchased from J. T. Baker and Sigma, respectively.

Arsenate resistance was assessed qualitatively on solid media by the radial streaking method [27]: Whatman GF/A filter disks (21 mm diameter) were placed in the middle of K medium plates containing 1 mM isopropyl-β-D-thiogalactopyranoside (IPTG; added to induce the expression of the cDNA genes from the *lac* promoter on the cloning vector), and 30 μl of 2 M Na_2_HAsO_4_ solution was placed on the disks and air-dried. Strains were streaked with sterile toothpicks outward from the disk, and their As^R^ was evaluated by the size of the zone of growth-inhibition next to the arsenate-containing filter disk compared to that seen with WC3110 *(ΔarsC)* carrying the empty cloning vector pTriplEx2. For a more quantitative determination of arsenate and arsenite resistance in liquid culture, we used the procedure of Mukhopadhyay et al. [12] with slight modifications: Cultures of strains carrying the *P. vittata* GST clones were grown overnight in LB + Ap and were subcultured at 1:100 dilution into K medium + 1 mM IPTG, plus different concentrations of Na_2_HAsO_4_ or NaAsO_2_ and grown with aeration. After 24.0 h of incubation at 37^°^, the cell density of the cultures was measured as light scattering in a Shimadzu UV-1700 spectrophotometer at 600 nm (OD_600_), with appropriate dilutions to correct for multiple scattering at OD_600_≥ 1.0 (where OD_600_ = 1 corresponds to ~1.5 x 10^9^ cells/ml). For each arsenate or arsenite concentration tested, the measurements were performed with three independent cultures.

### *Pteris vittata* cDNA library construction and screening

RNA was purified from *P. vittata* gametophytes grown for 6 weeks in liquid culture [28] containing 1 mM KH2AsO4. A cDNA library was synthesized using BD Biosciences Clonetech SMART^®^ cDNA library Construction kit (User Manual PT3000-1 [PR15738]) and cloned between the SfiIA and *Sfi*IB sites of plasmid pTriplEx2 (Clonetech). The ligation mixture was electroporated into *E. coli* strain DH10B [29] and transformants selected on LB Ap plates. Plasmid DNA was isolated from a pool of ~7x10^6^ colonies and electroporated into WC3110 (ΔarsC). The cells were allowed to recover in 1 ml LB medium for one hour at 37^°^C with aeration. Aliquots of 100 μl of the transformation mixtures were inoculated on K medium plates containing 6mM Na_2_AsO_4_·7H_2_O and 1 mM IPTG. After two days, the plates were examined for the appearance of As^R^ colonies. Spontaneous As^R^ mutants arose in the untransformed control cells at a frequency of ~2 per 10^7^ cells; the frequency of such derivatives was generally 2-to 10-fold higher in the cultures transformed with the library. For plates that contained fewer than ~50 As^R^ transformed colonies, the strains were single colony isolated on K medium plates containing 6 mM Na_2_AsO_4_·7H_2_0, and then tested for Ap resistance (Ap^R^) on LB plates, confirming the presence of the cloning vector (as opposed to a spontaneous mutation to As^R^ by the host). Plates that contained more than 50 As^R^ colonies were replica-plated to LB Ap plates, and colonies that were stably Ap^R^ were single colony isolated on K medium plates containing 6 mM arsenate. Plasmids were isolated from the transformants that were confirmed to be both As^R^ and Ap^R^ and were retransformed into WC3110, selecting Ap^R^. The derivatives obtained in the latter step were tested for As^R^ by radial streaking. Plasmids were isolated from the As^R^ strains and the sequences of the cDNA inserts were determined at Purdue Genomics Core Facility. The sequences were analyzed with the blastn and blastp algorithms (http://blast.ncbi.nlm.nih.gov/Blast.cgi). The sequences of the three *P. vittata* ST cDNA clones *pPvGST35, pPvGST41,* and *pPvGST57* that we obtained were entered into GenBank under accessions numbers EU259318, EU259319, and EU259320, respectively.

### Speciation of arsenic in cell cultures

Strains KC1749 (pTriplEx2), KC1761 *(pPvGST35)* and KC1763 *(pPGSTv41)* were grown LB + Ap, 0.03 ml were inoculated into 3 ml of K medium containing 10 mM glucose + 0.1 mM Na_2_HAsO_4_ + Ap + 1 mM IPTG and incubated with aeration for 24 h. At the end of the growth, the cell density was determined by light scattering at 600 nm (OD_600_). The cells from 1.5 ml samples were sedimented by a microfuge, 0.5 ml of the supernatant culture media were mixed with 0. 5 ml of a buffer containing 13.36 mM NH4NO3, 13.36 mM NH4H2PO4, and 10 mM EDTA (pH 6.2) and flash-frozen in dry ice ethanol. EDTA was used in the extraction buffer as it has been shown to preserve the inorganic arsenic species [30]. The As(III) and As(V) content of the samples from 5 independent cultures for each strain were determined as described by the method of Williams et al. [31].

## Results

### Isolation of cDNA clones from *P. vittata* that confer arsenate-resistance in *E. coli*

Plasmids *pPvGST35, pPvGST41,* and *pPvGST57* were isolated by introducing a *P. vittata* cDNA expression library into the *ΔarsC* (arsenate reductase) mutant *E. coli* WC3110 and selecting derivatives that could grow in K medium containing 6 mM Na_2_AsO_4_. The arsenate resistance (As^R^) phenotype of these transformants is illustrated in Figure 1: The three plasmids carrying the *P. vittata* cDNA clones conferred increased As^R^ on this host strain compared to the control carrying the empty vector pTriplex2. The phenotype conferred by these three plasmids was evaluated in a more quantitative form in Figure 2, which shows that in liquid medium, the plasmids conferred very similar increases in As^R^ above that seen with the control strain carrying the empty cloning vector. WC3110 derivatives expressing the *P. vittata* GST clones had an exponential doubling time of 1.6 h in K medium +10 mM Na_2_AsO_4_, whereas the empty vector control had a doubling time of >12 h (data not shown).

**Figure 1.**
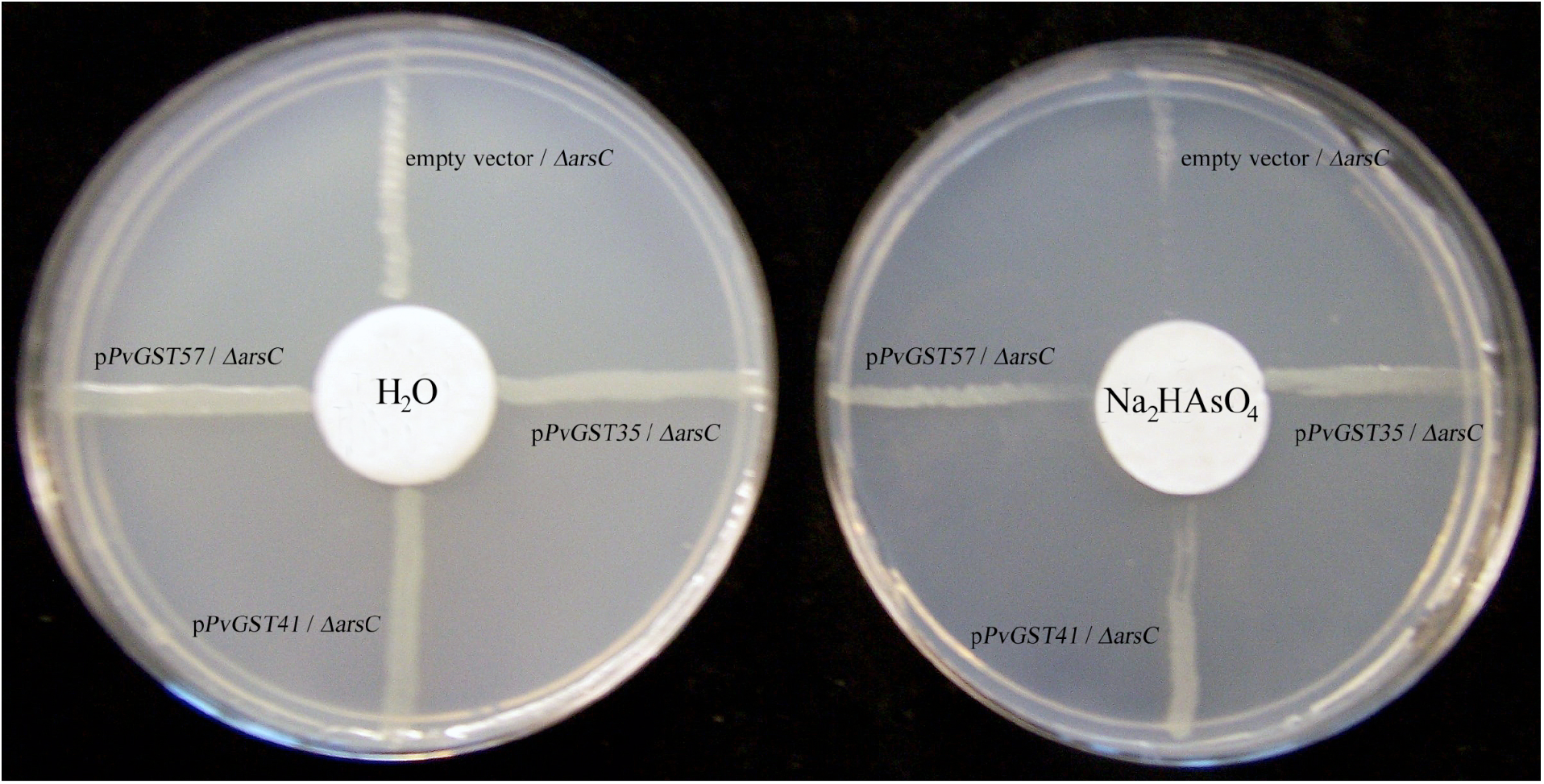
Qualitative demonstration of the arsenate resistance conferred by the *P. vittata* GST clones in *E. coli.* Derivatives of strain WC3110 *(ΔarsC)* carrying the *P. vittata* clones *PVGST35, PVGST41,* and *Pv57* were tested for arsenate resistance on solid media by radial streaking, as described in Materials and Methods. The filter disk placed on the Petri dish on left was spotted with 30 of μl H2O and the one on the right was spotted with 30 μl of 1.8 M Na_2_HAsO_4_.

**Figure 2.**
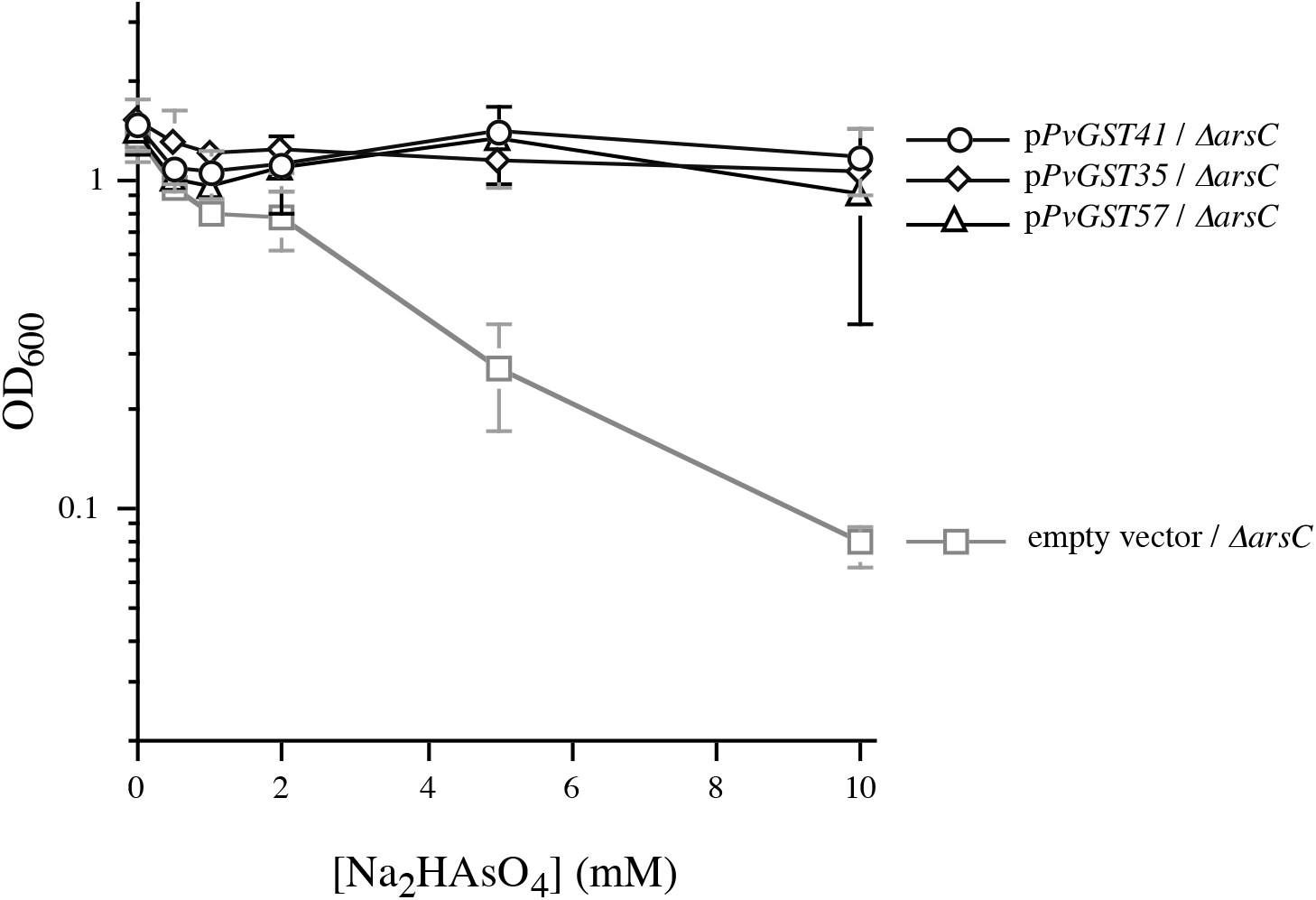
Arsenate resistance conferred by the *P. vittata* GST clones on *E. coli* in liquid cultures. Resistance to the indicated concentrations of Na_2_HAsO_4_ was determined in liquid cultures, as described in Materials and Methods. Strains used were derivatives of WC3110 (*ΔarsC*) carrying pTriplEx2 (empty vector), pTriplEx2 containing *P. vittata* GST1 (p*PvGST35* and *pPvGST57),* or *P. vittata* GST2 *(pPvGST41).* The results are the averages ± standard deviation (indicated by error bars) of data obtained with 3 independent cultures of each strain.

### The cDNA clones conferring arsenate resistance encode GST-family proteins

DNA sequence analysis revealed that the three plasmids conferring As^R^ contain a 217-codon open reading frame whose deduced protein product shows high similarity to GSTs from higher plants. Plasmids *pPvGST35* and *pPvGST57* are identical in their nucleotide sequences in the central protein-coding region (Figure 3); the gene contained in these two plasmids was designated *PVGST1.* These two plasmids exhibited slight sequence differences at the 5’ and 3’ ends of the inserts, which could have been due to the fact that they were from different mRNAs, or differences could have been introduced during cloning. In any case, the fact that the plasmids are different indicates that they are independent isolates. The insert in *pPGSTv41* shows a 13% nucleotide sequence difference from the inserts in *pPvGST35* and *pPvGST57,* resulting in 23-amino acid difference in the predicted open reading frames (Figure 3). Consequently, the open reading frame in *pPvGST41,* which is designated *PVGST2,* specifies a GST that is different from *PVGST1.* The expression vector pTriplEx2 contains translation start signals for 5’ portions of the *ompA* and *lacZ* genes in different reading frames, followed by a run of T residues that provide a site for transcriptional frameshifting. Consequently, cloned inserts may be translated in all three reading frames as N-terminal fusion proteins with OmpA or LacZ peptides. DNA sequencing revealed that the inserts in *pPvGST35* and *pPvGST57* contain a TAG termination codon 24 bases upstream of the ATG initiation codon for the GST, indicating that these two gene products are not made as fusion proteins. However, there is no analogous in-frame termination codon upstream of the GST sequence in *pPvGST41,* and so the latter may be translated as a fusion product with an N-terminal peptide from LacZ or OpmA.

**Figure 3.**
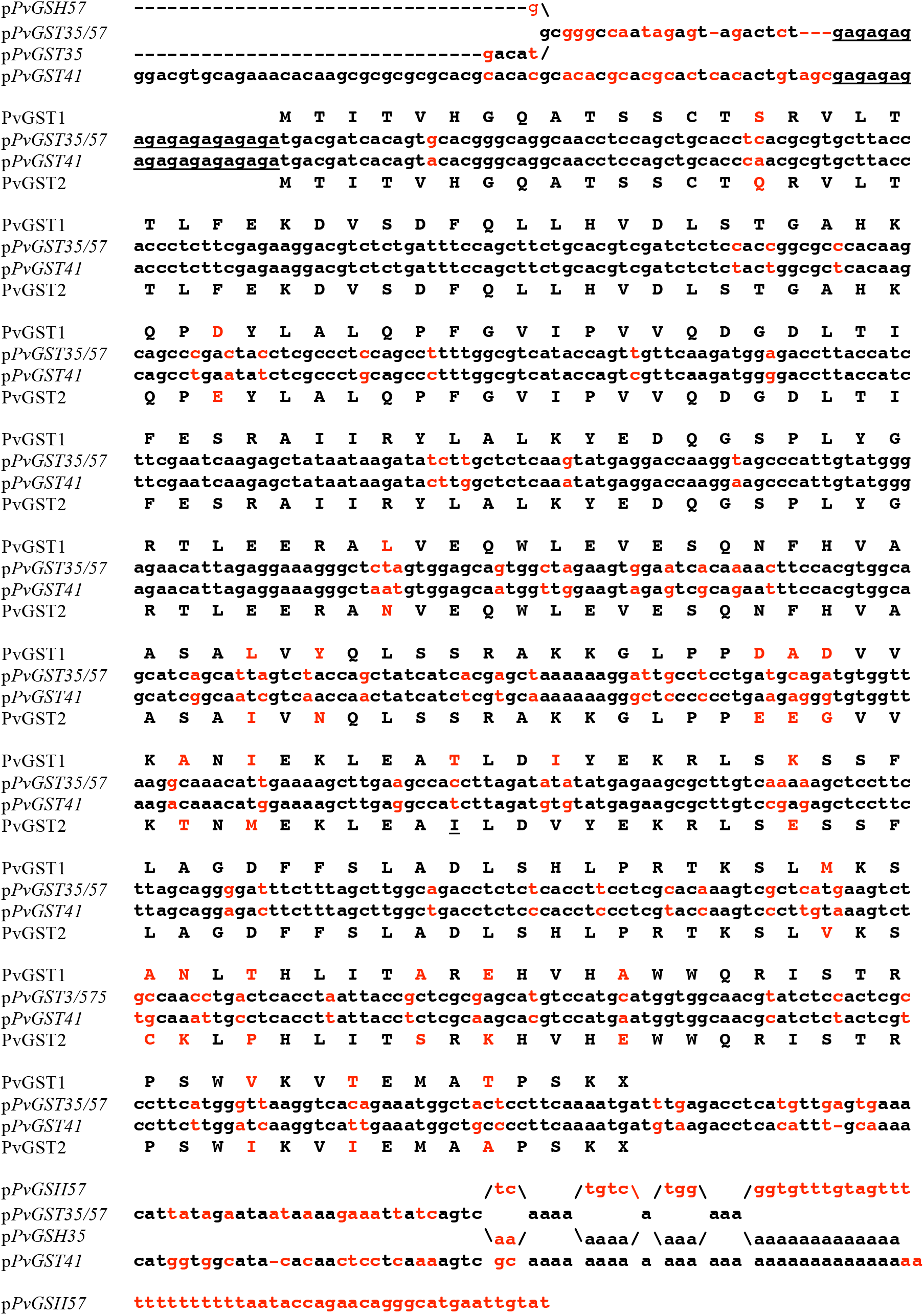
Sequences of the *P. vittata* GST cDNA clones. The nucleotide and predicted amino acid sequences of the*pPvGST35,pPvGST57,* and*pPvGST41* cDNA clones are shown with lower case and upper case letters, respectively. Nucleotide sequences that are common to p*PvGST35* and *pPvGST57* are indicated as “*pPvGST35/57*where the two clones differ, the nucleotide sequences are offset above (for *pPvGST57)* and below (for *pPvGST35)* the common sequences. The predicted amino acid sequences of PVGST1 and PVGST2 are shown by capital letters above and below the nucleotide sequences for *pPvGST35/57* and *pPvGST41,* respectively. The nucleotide and amino acid sequences that are common to p*PvGST35* and p*PvGST41* are shown in black letters, and sequences that are different between the two clones are shown in red. The GA repeats before the translation start sites are underlined.

GSTs of higher plants have been assigned into four classes based on their amino acid sequences: phi, tau, theta, and zeta; representatives of the former two classes are also found in bacteria and members of the latter two classes are also present in animals [32]. Figure 4A shows a multiple alignment of the two *P. vittata* genes against four of the most closely related enzymes from other plant species and one representative of each of the four classes of GSTs from *A. thaliana.* This analysis indicates that the *P. vittata* enzymes cluster together and with other members of the phi-class of plant GSTs (Figure 4B). One of the diagnostic features of the phi class of GSTs is that their GSH-binding site contains the amino acid triplet ESR [32], which is present in the *P. vittata* clones (Figure 4A).

**Figure 4.**
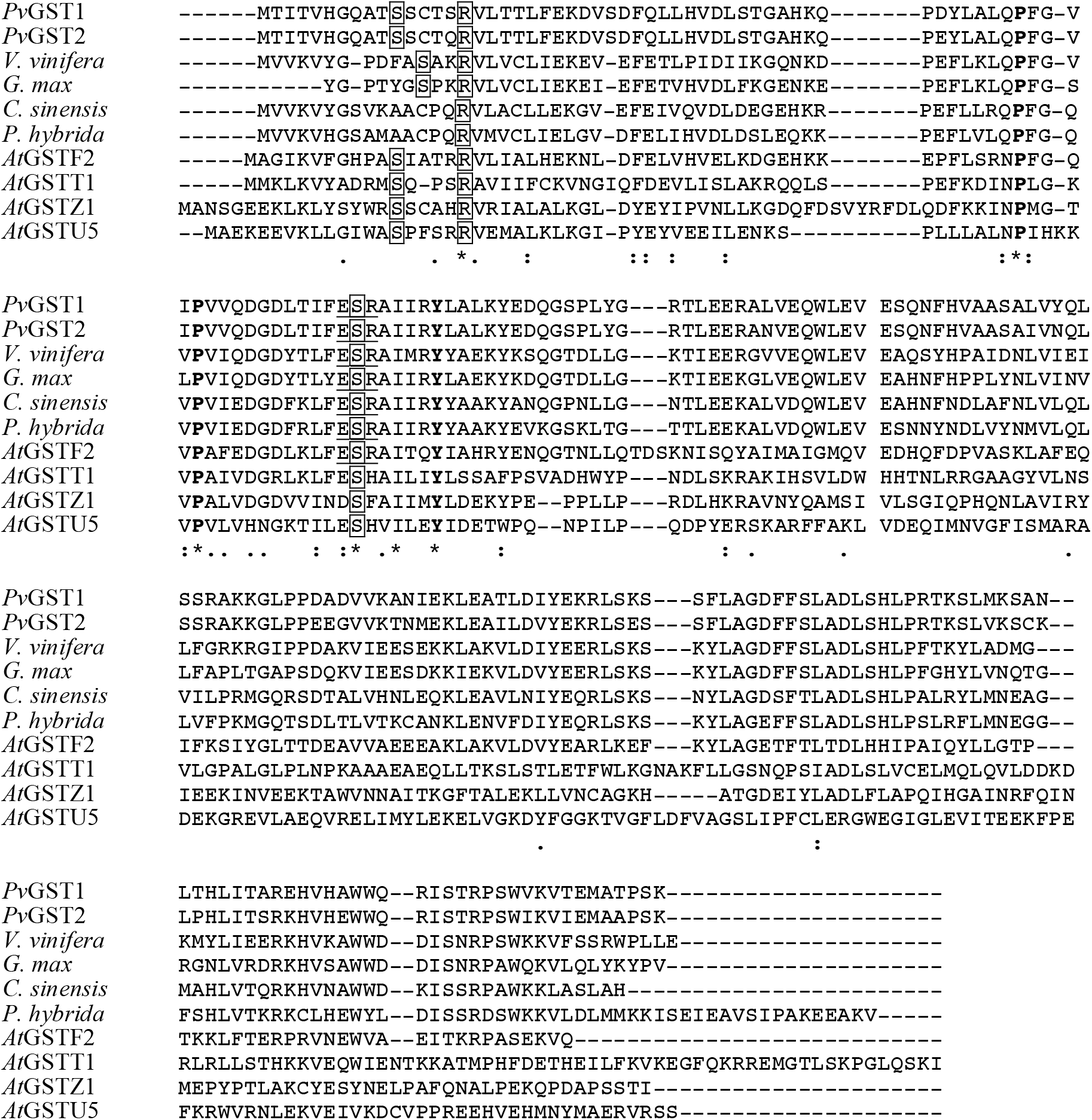
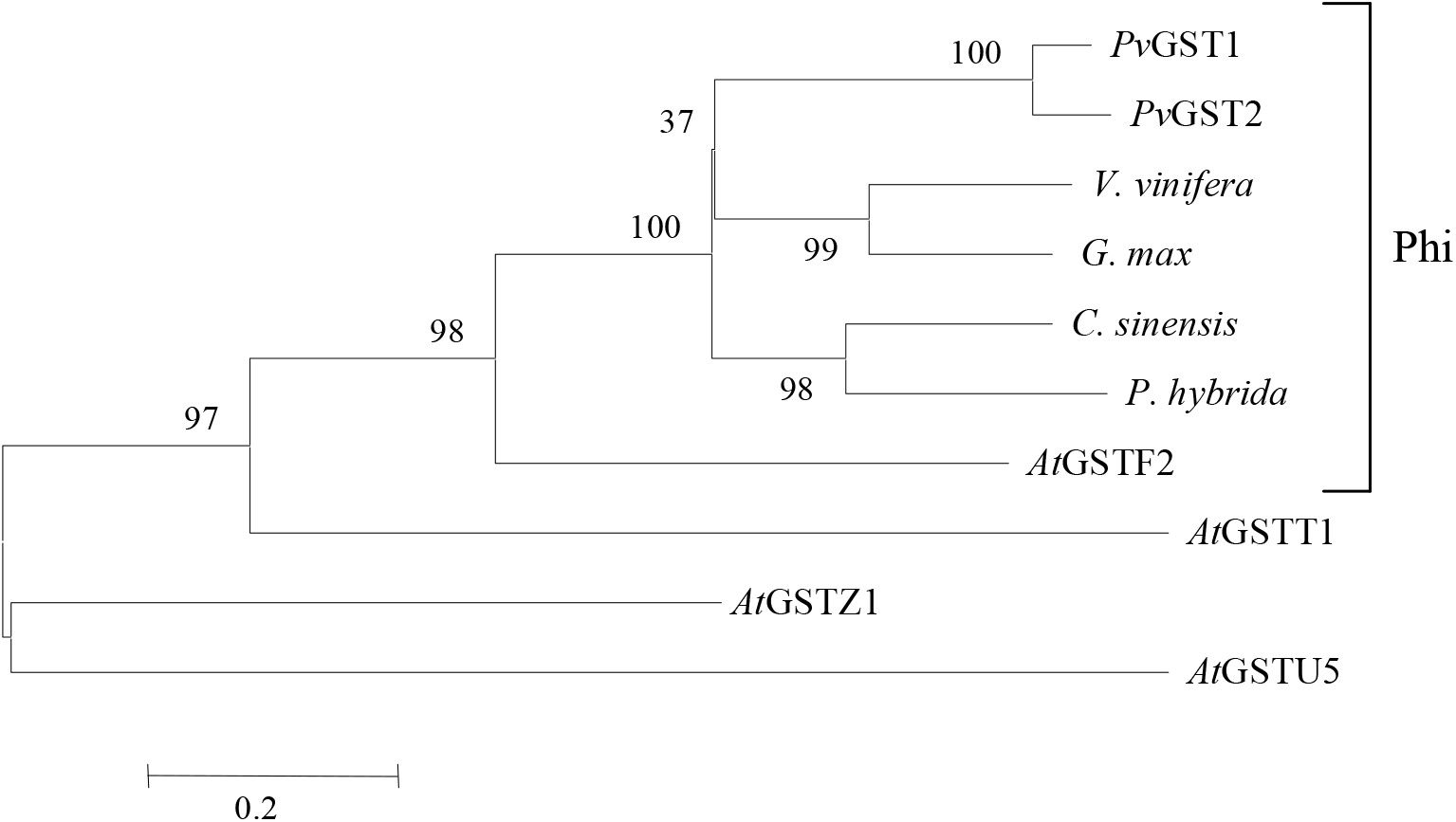
Sequence relationship of the *P. vittata* GSTs with orthologs from other plants. Panel A: Alignment of the amino acid sequences of *PVGST1* and *PVGST2* with the four plant GSTs that showed the highest similarity to *PVGST1* (blastp Expect values between 1e-60 and 2e-62), and with one representative of each of the four classes of GSTs from *A. thaliana.* Multiple sequence alignment was performed with the MultAlin algorithm (http://multalin.toulouse.inra.fr/multalin) [47]. The amino acids that are in boxes are conserved in most GSTs and have been shown to constitute part of the catalytic site, and the amino acids shown in **bold** letters have been found to be conserved in all *A. thaliana* GSTs [32]. The underlined <u>ESR</u> sequence is diagnostic of the phi-class of GSTs and constitutes part of the GSH-binding site [32]. The characters “*”, “:”, and “.” below the comparisons indicate fully conserved, strongly conserved and weakly conserved residues, respectively, as defined in the instructions for the CLUTALW algorithm (http://align.genome.jp/). The sequences used were [accession numbers in brackets]: *P. patens*, moss *Physcomitrella patens subsp. patens* hypothetical protein GST N family [XP_001764210.1]; *P. sitchensis,* Sitka spruce *Picea sitchensis* hypothetical protein GST C family [ABK23015.1]); *C. sinensis,* navel orange *Citrus sinensis* cultivar Tarocco glutathione S-transferase N family [ABA42223]; *C. argyrophylla*, conifer yin shan *Cathaya argyrophylla* phi class glutathione transferase [ACL37469.1]; AtGSTF2, *A. thaliana* phi-class glutathione S-transferase F2 [NM_116486.2]; ATGSTT1, *A. thaliana* theta-class glutathione S-transferase T1 [NM_123486.3]; AtGSTZ1, *A. thaliana* zeta-class glutathione S-transferase Z1 [NM_201671.1]; AtGSTU5, *A. thaliana* tau-class glutathione S-transferase U5 [NM_128499.3]. **Panel B**: The dendogram was generated from the multiple alignment in Panel with the MultAlin algorithm.

The *P. vittata* GSTs do not show significant similarity (blastp Expect value > 0.019) to the three types of arsenate reductases represented by the *E. coli*, *S. aureus*, and the yeast/*P. vittata* enzymes. Genes encoding highly similar phi class GST can be found in the *Myxococcales* bacteria *Sorangium cellulosum* ‘So ce 56’ and *Myxococcus xanthus* DK 1622 (blastp similarity to *PVGST1* Expect value = 5e-43 and 6e-28, respectively) and in over 100 other bacteria with blastp Expect values between 1e-18 and 1e-13. The function of these bacterial enzymes has not yet been determined.

### Nucleotide sequence features of the *P. vittata* GST clones

An interesting feature of the three *P. vittata* GST clones is that they contain 10 repeats of the dinucleotide GA immediately upstream and overlapping the ATG start (Figure 3). This motif is not present in the most closely related plant GSTs genes, including the one from the moss *Physcomitrella patens.* However, a cDNA glutaredoxin clone from *P. vittata* [23] also contains 12 GA repeats seven nucleotides upstream of the translation start site (see sequence under GenBank accession number EF052272). Whether these repeats are associated with the expression of arsenic tolerance-related genes in *P. vittata* or whether they have some other function in gene expression in this fern remains to be explored. There are precedents for strings of GA repeats upstream of genes in other eukaryotic systems. In *Drosophila melanogaster*, the so-called GAGA elements have been recognized as the binding sites of a chromatin remodeling factor that increases transcription from the *hsp70* promoter [33 and references therein]. A GAGA element has been found between the TATA box and the transcription start sites of the *Gsa1* gene of *Glycine max,* which has been suggested to be involved in the transcriptional control of the gene. Repeats of GA, or its complement CT, have also been found downstream of viral promoters, and it has been suggested that they play a positive role in transcriptional activation [34,35].

### Phenotypic characterization of the *P. vittata* GST clones in *E. coli*

We tested whether the *P. vittata* GST clones can confer increased As^R^ in an arsenate reductase proficient *(arsC+)* host (Figure 5). Our results reproduce the finding of Mukhopadhyay et al. [12] that loss of the *E. coli* arsenate reductase greatly increases sensitivity to arsenate (compare arsC+/empty vector with ΔarsC/empty vector). Although the *P. vittata* clones substantially elevate As^R^ in the *ΔarsC* mutant, they resulted only in a marginal, statistically insignificant increase in the already high As^R^ of the *arsC+* strain. These results indicate that although the *P. vittata* GSTs can functionally substitute for the *E. coli* arsenate reductase in the *ΔarsC* mutant, they do not provide improved resistance against this toxic chemical above the level conferred by the endogenous enzyme.

**Figure 5.**
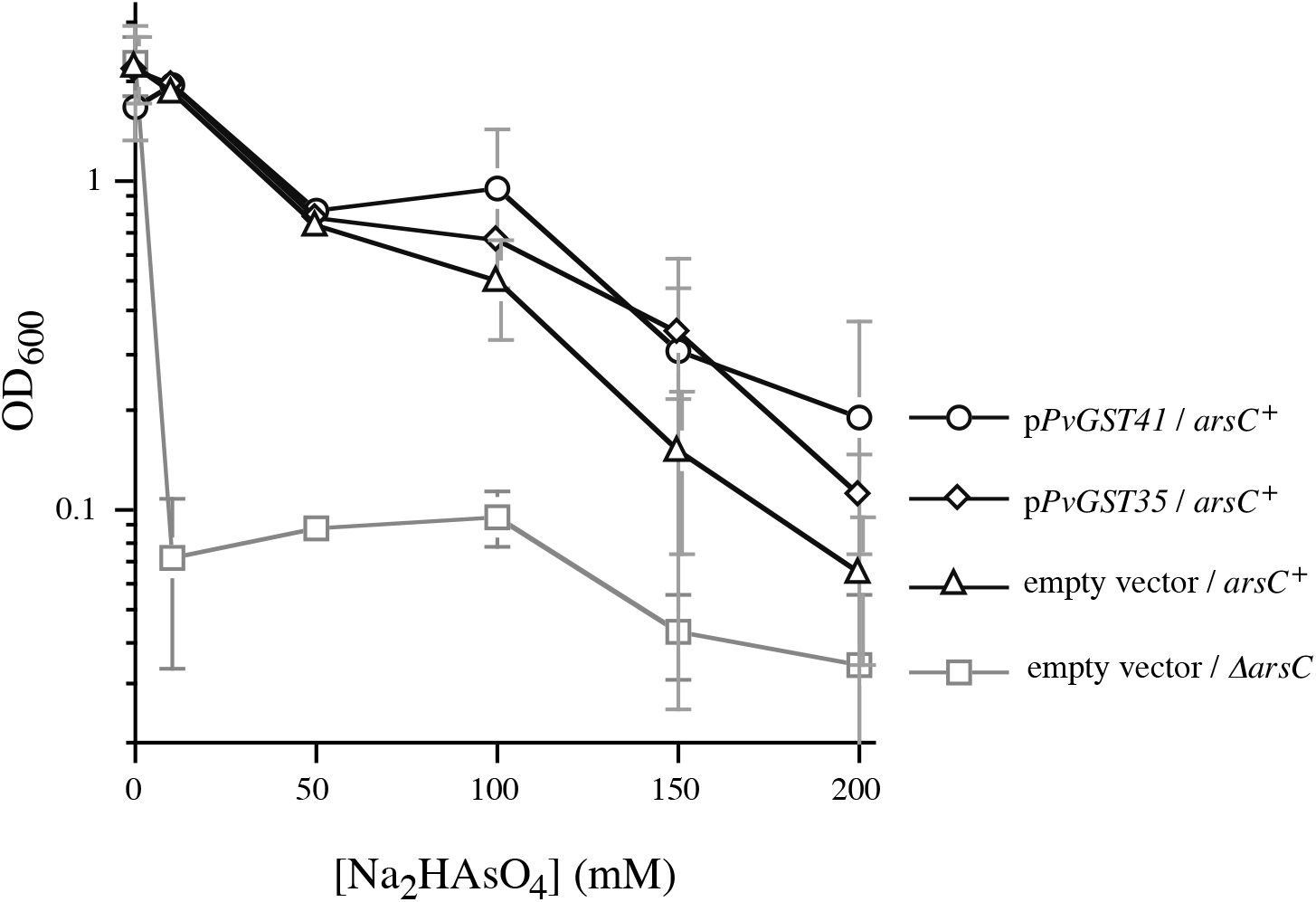
Arsenate resistance is not enhanced by *P. vittata* GST cDNAs in wild-type *E. coli.* Resistance was determined in liquid cultures as described in Materials and Methods in the arsenate reductase proficient *(arsC+)* W3110 strain carrying pTriplEx2 (empty vector), pTriplEx2 containing *P. vittata* GST1 *(pPvGST35),* or *P. vittata* GST2 *(pPvGST41).* Also included are data for WC3110 *(ΔarsC)* carrying the empty vector pTriplEx2. The results are the averages ± standard deviation (indicated by error bars) of data obtained with 3 independent cultures of each strain.

The sequence similarity of our clones to known GSTs suggests that the *P. vittata* enzymes probably use glutathione as a substrate. To test this, we determined whether blocking glutathione synthesis with a *gshA* (*γ*-glutamyl-cysteine synthetase) mutation would diminish the arsenate resistance conferred by these plasmids. The *gshA ΔarsC* double mutant carrying the empty cloning vector was considerably more sensitive to arsenate in the absence of GSH than the *ΔarsC gshA+* counterpart (Figure 6A). More importantly, the *gshA* mutation cancelled the ability of the *P. vittata* clones to confer arsenate resistance in the absence of glutathione. Supplementation with GSH, which can be taken up by *E. coli* [36], restored the ability of the clones to confer As^R^ in the *gshA* mutant background (Figure 6B). These results suggest that the *P. vittata* GSTs use GSH as substrate for the detoxification of arsenate.

**Figure 6.**
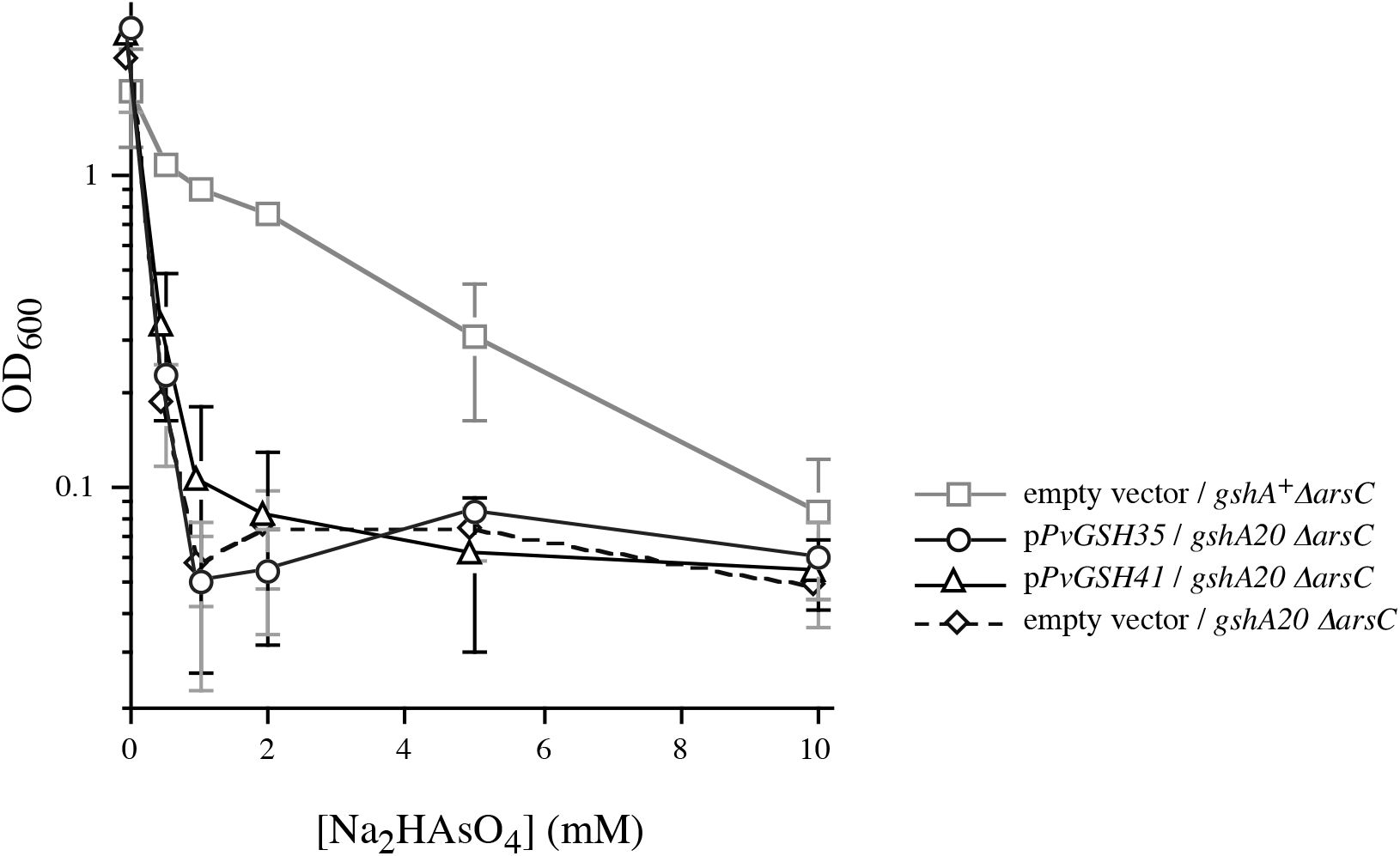
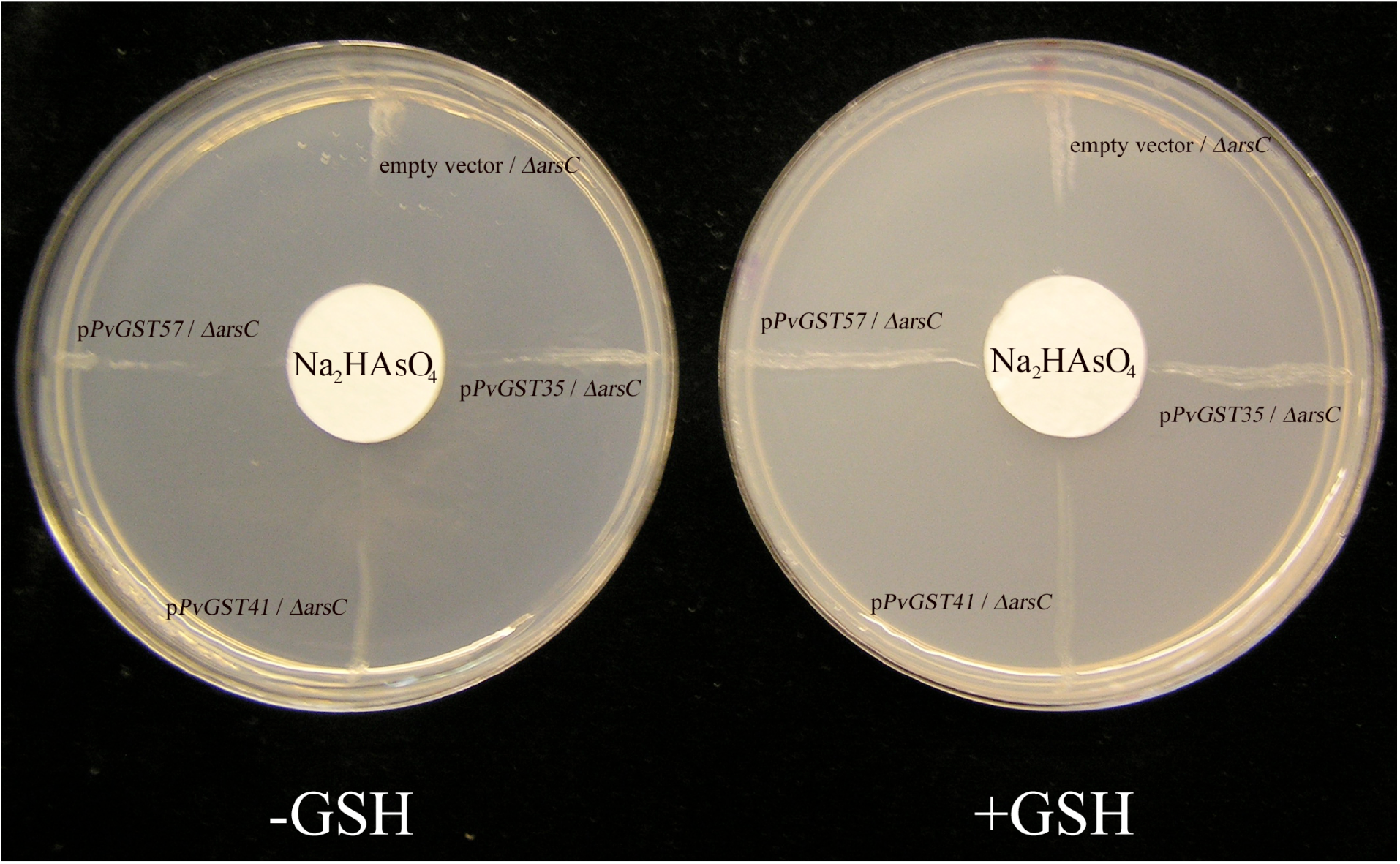
Glutathione is required to confer arsenate resistance by the *P. vittata* GST cDNAs. **Panel A**: Arsenate resistance was determined in a *gshA20 ΔarsC* double mutant in liquid cultures in the absence of reduced glutathione, as described in Materials and Methods. The strains carried pTriplEx2 (empty vector), *pPvGST35,* and *pPvGST41.* Also included is a control with strain WC3110 *(gshA+ ΔarsC)* carrying pTriplEx2 (empty vector). The results are the averages ± standard deviation (indicated by error bars) of data obtained with 3 independent cultures of each strain. **Panel B**: Arsenate resistance of the *gshA20 ΔarsC* strains carrying the indicated plasmids was determined on solid media, as described in the Materials and Methods. The Petri dish on the left contained K medium and the one on the right contained K medium augmented with 1 mM GSH, and the filter disks on both plates were spotted with 30 μl of 1.8 M Na_2_AsO_4_.

Finally, we examined whether the *P. vittata* GSTs could confer resistance to arsenite and found that that the clones were unable to do so either in the *ΔarsC* mutant (Figure 7) or in the *arsC^+^* background (data not shown), indicating that the *P. vittata* GSTs can detoxify arsenate but not arsenite.

**Figure 7.**
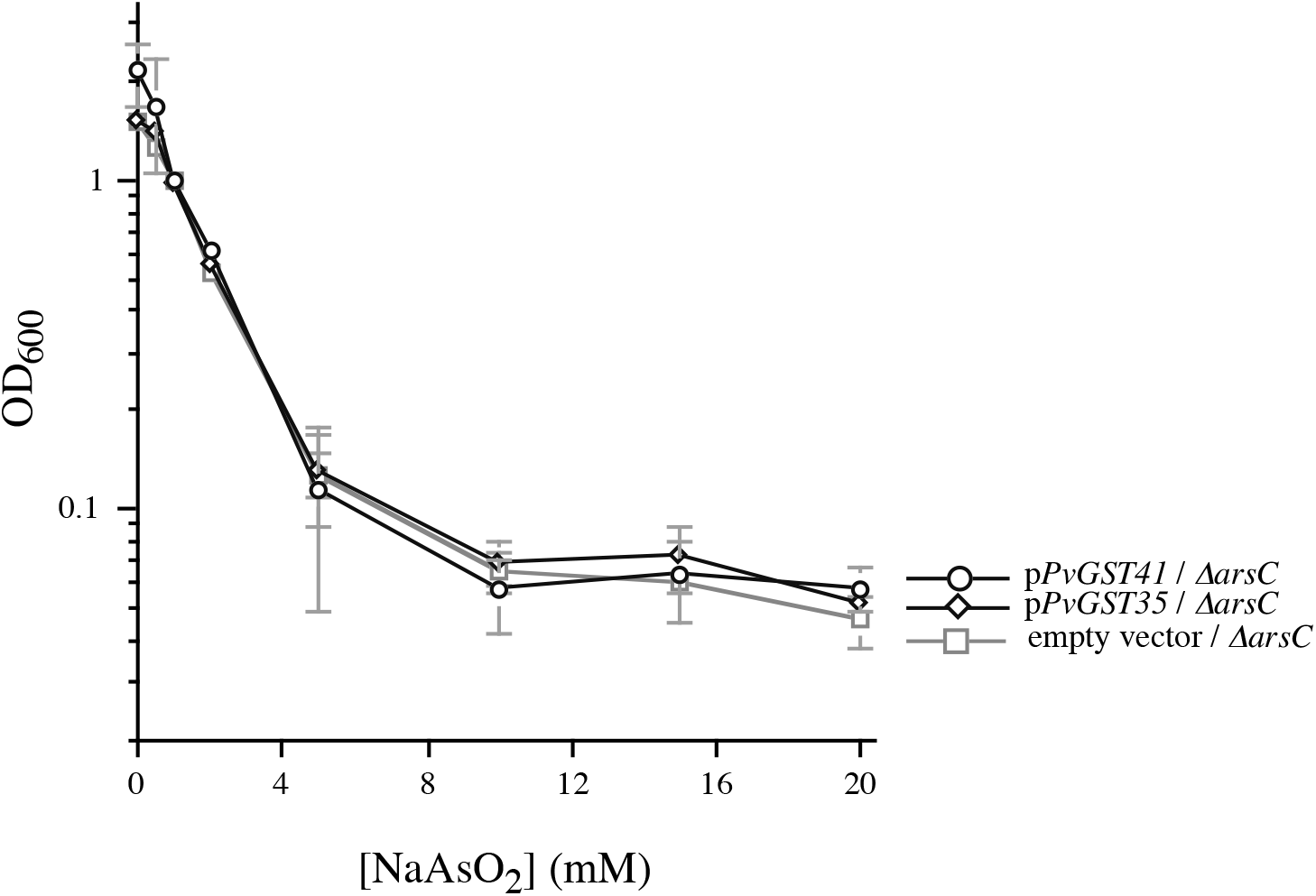
The *P. vittata* GST clones do not increase resistance to arsenite. Resistance to the indicated concentrations of NaAsO_2_ was determined in liquid cultures, as described in Materials and Methods. Strains tested were derivatives of WC3110 *(ΔarsC)* carrying pTriplEx2 (empty vector), pTriplEx2 containing *P. vittata* GST1 (p*PvGST35*), or *P. vittata* GST2 (p*PvGST41*). The results are the averages ± standard deviation (indicated by error bars) of data obtained with 3 independent cultures of each strain.

We carried out GST activity assays of the *E. coli* strains carrying the *P. vittata* clones using 1-chloro-2,4-dinitrobenzene (CDNB) as substrate, which has been used to assay phi class GSTs from plants [32,37]. However, we were not able to detect statistically significant GST activity in the *E. coli* strains carrying the *P. vittata* clones (data not shown), indicating that the *P. vittata* GSTs are unusual among the phi class enzymes in that they have no detectable activity with CDNB as substrate.

### The *P. vittata* GSTs promote the reduction of arsenate in *E. coli*

In order to probe the mechanism by which the *P. vittata* GST clones confer increased arsenate resistance, we carried out two experiments to determine whether this phenotype involves the reduction of this toxic compound. In one of these experiments, we tested the effect of a glutathione reductase (gor) mutation on the ability of the *P. vittata* GSTs to impart increased arsenate resistance. The result was that the ability of the GST plasmids *pPvGST35* and *pPvGST41* to confer increased arsenate resistance was compromised by the *Δgor::cat* mutation (Figure 8). This result suggests that arsenate detoxification involves the oxidation of GSH, and glutathione reductase is required to regenerate this reduced sulfhydro compound.

**Figure 8.**
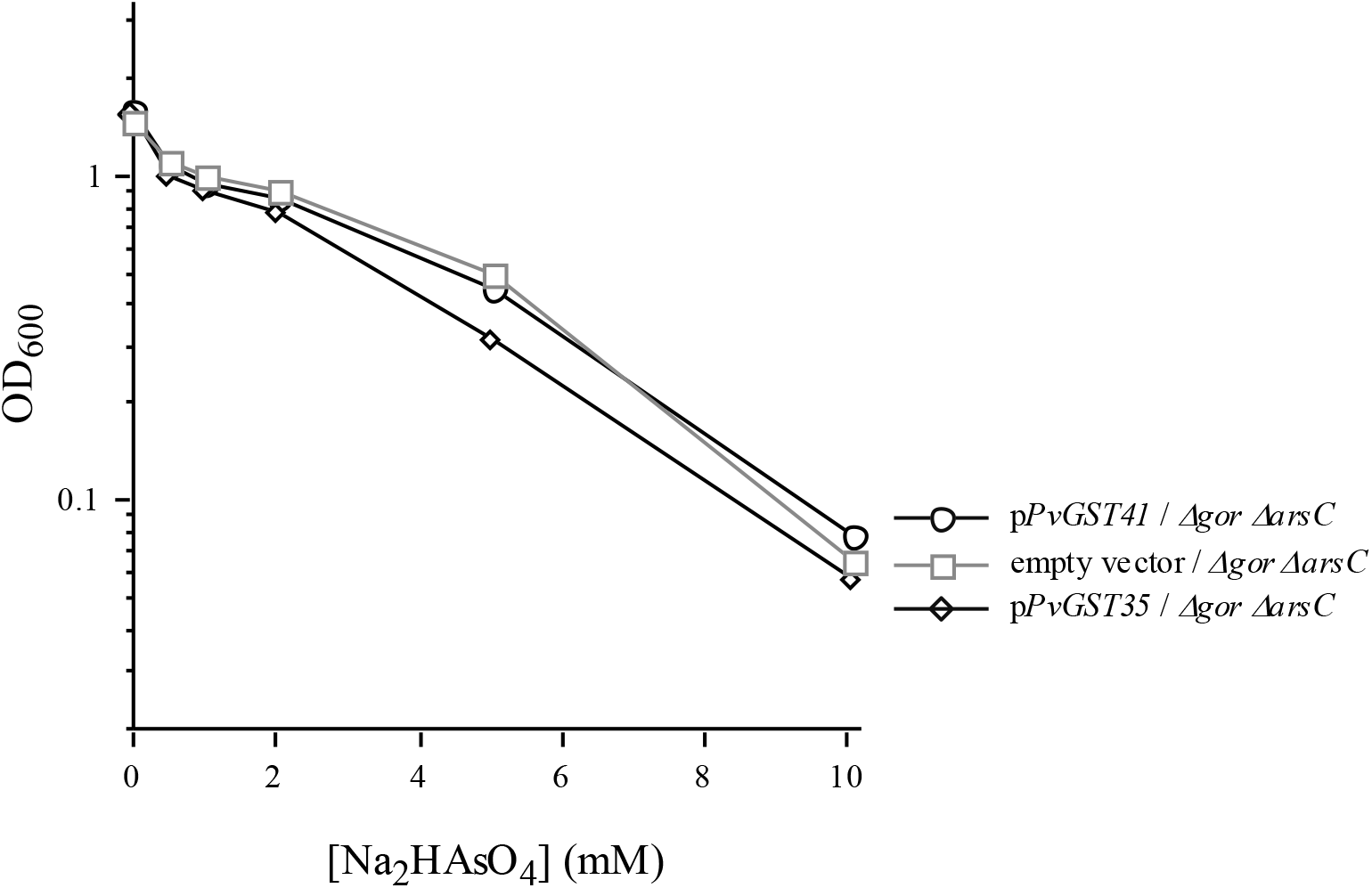
Glutathione reductase is required for the ability of the *P. vittata* GST clones to confer arsenate resistance. Resistance to the indicated concentrations of Na_2_HAsO_4_ was determined in liquid cultures, as described in Materials and Methods. Strains tested were derivatives of KC1852 *(ΔarsC::kan Δgor::cat)* carrying pTriplEx2 (empty vector), pTriplEx2 containing *P. vittata* GST1 (p*PvGST35*), or *P. vittata* GST2 (p*PvGST41*). The results are the averages ± standard deviation (indicated by error bars) of data obtained with 3 independent cultures of each strain.

We also tested whether the *P. vittata* GSTs increases the formation of As(III) from arsenate. For this experiment, we grew derivatives of strain WC3110 *(ΔarsC)* carrying the empty cloning vector or plasmids p*PvGST35* and p*PvGST41* in medium containing an initial 1 mM Na_2_HAsO_4_, and after 24 h of growth, we quantified the proportions of arsenic as As(III) and As(V) in the culture medium. The final biomass density of the cultures was OD_600_ = 0.96 ± 0.05 for the strain carrying the empty cloning vector, and OD_600_ = 1.71 ± 0.10 and 1.82 ± 0.15 for the strains carrying p*PvGST35* and p*PvGST41*, respectively, confirming the ability of the GST plasmids to confer increased arsenate resistance. The cells were sedimented by centrifugation, and the amounts of arsenic in the medium as As(III) and As(V) were determined as described in Materials and Methods. The results are shown in Table 1. For strain KC1749, which carried the empty cloning vector, approximately 30% of the input arsenic was converted to As(III), with the balance remaining as As(V). It is not clear whether the reduction of arsenate in this culture was due to an enzymatic process or to some spontaneous reaction with GSH or some other sulfhydro compound. However, in the cultures of the strains carrying p*PvGST35* and p*PvGST41*, the proportion of arsenic that was recovered as As(III) increased to approximately 60% and 69%. This result indicates that the *P. vittata* GSTs promoted the formation of an As(III) species, suggesting that the increased arsenate resistance conferred by these enzymes involves the reduction of arsenate.

**Table 1.** Conversion of As(V) to As(III) by strains synthesizing the *P. vittata* GSTs

| Strain | As(III)<br>($\mu\text{mol} / \text{ml culture}$) | As(V)<br>($\mu\text{mol} / \text{ml culture}$) | As(III)<br>% of total As |
| --- | --- | --- | --- |
| KC1749<br>(empty vector<br>control) | $0.24 \pm 0.04$ | $0.55 \pm 0.09$ | $30.3 \pm 0.5$ |
| KC1761<br>(pPvGST35) | $0.51 \pm 0.08$ | $0.35 \pm 0.08$ | $59.6 \pm 3.3$ |
| KC1763<br>(pPvGST41) | $0.57 \pm 0.07$ | $0.26 \pm 0.04$ | $68.7 \pm 0.7$ |

The strains were grown in medium containing $1.0 \mu\text{mol} / \text{ml Na}_2\text{HAsO}_4$ and the speciation of arsenic as As(III) and As(V) was determined after 24 h of growth, as described in Materials and Methods. Results are averages ± standard deviation of data obtained with 5 independent cultures of each strain.

## Discussion

In this work, we isolated three independent clones from a *P. vittata* cDNA library that imparted increased arsenate resistance to an *arsC* mutant *E. coli.* Sequence analysis indicated that these clones encode proteins that have high sequence similarity to the phi family of higher plant GSTs. GSTs have diverse functions related to the detoxification of harmful substances [32]. Some have glutathione dependent peroxidase or isomerase activities [38,39]. Others are involved in detoxification of xenobiotics by glutathione conjugation. It has been suggested that some GSTs function as ligands or carrier proteins for a variety of molecules [40]. They have also been implicated in signal transduction pathways [41] and in responses to a wide range of stresses [32].

Our phenotypic analysis in a *ΔarsC* mutant E*. coli* indicated that the *P. vittata* GSTs require GSH for arsenate resistance. The ArsC, Acr2p, and PvACR2 arsenate reductases of E. *coli,* yeast, and *P. vittata* use GSH as electron donor and produce oxidized glutathione (GSSG) [12,13,42]. Arsenate reduction in *E. coli* and yeast requires the participation of a glutaredoxin, which regenerates GSH from GSSG. However, the human monomethylarsonic acid reductase, which is a member of the omega family of mammalian GSTs [43], can carry out the GSH-dependent reduction of methylarsonate to methylarsenite without the direct involvement of a glutaredoxin [44]. It is possible that the *P. vittata* GSTs may be able to catalyze the analogous reduction of arsenate to arsenite.

Another type of mammalian GSTs, which belong to the omega subfamily, can detoxify arsenite by forming an arsenite-(GSH)3 complex that is excreted from the cells [45,46]. Our observation that the *P. vittata* GSTs do not provide increased resistance to arsenite suggests that they do not operate by this mechanism. It is possible that the *P. vittata* GSTs might facilitate the formation of an arsenate-GSH adduct that could be expelled from *E. coli.* However, the latter mechanism would necessitate the stoichiometric consumption of at least one molecule of GSH for one molecule of arsenate that is detoxified. In view of the fact that our GST cDNA clones could provide resistance to at least 10 mM arsenate in a medium that contains only 10 mM glucose as carbon source, we feel that this mechanism is unlikely. The requirement for glutathione reductase also supports the conclusion that GSH has a catalytic rather than a stoichiometric role.

Arsenate resistance in the hyperaccumulator *P. vittata* is thought to involve reduction of As(V) to As(III) and transport into the vacuoles in fronds and gametophytes. An arsenate reductase has been cloned from *P. vittata* [13], and it has been recently reported that a triosephosphate isomerase [22] and glutaredoxin [23] from this organism can increase the arsenic resistance in *E. coli.* Our observations that the *P. vittata* GSTs promote the GSH-dependent conversion of As(V) to As(III) in the *arsC* mutant *E. coli* suggest that these proteins participate in an arsenate reduction pathway, which also involves the bacterial glutathione reductase. However, we do not have any direct evidence that the *P. vittata* GSTs by themselves have arsenate reductase activity and cannot rule out other possible mechanisms, such as that these enzymes might detoxify arsenate by promoting the formation of a GSH complex. It will be necessary to purify these enzymes from *E coli* and establish whether they have one of the above enzymatic activities in vitro. Finally, although the fact that the GSTs conferred increased arsenate tolerance in *E. coli* makes it attractive to hypothesize that these proteins are likely to contribute to the arsenic tolerance or hyperaccumulation in *P. vittata*, it will be necessary to confirm the roles of these enzymes in arsenic metabolism in the ferns.

## Acknowledgments

We thank A. Murphy for helpful discussions, and N. Csonka for help with the photography and image processing.

## Author Contributions

Conceived and guided the project: DES LNC. Isolated *P. Vittata* GST cDNA clones in *E. coli* and carried out initial phenotypic characterization: AAK. Constructed the *P. vittata* cDNA expression library: DRE. Carried out the arsenic speciation and related data analysis: GJN and AAM. Performed the GST assays: XH. Wrote the manuscript: LNC. All authors read and approved the final manuscript.

